# Chromatin Landscapes and Genetic Risk For Juvenile Idiopathic Arthritis

**DOI:** 10.1101/075721

**Authors:** Lisha Zhu, Kaiyu Jiang, Karstin Webber, Laiping Wong, Tao Liu, Yanmin Chen, James N. Jarvis

## Abstract

Juvenile idiopathic arthritis (JIA) is considered to be an autoimmune disease mediated by interactions between genes and the environment. To gain a better understanding of the cellular basis for genetic risk, we studied known JIA genetic risk loci, the majority of which are located in non-coding regions, in human neutrophils and CD4 primary T cells to identify genes and functional elements located within those risk loci. We analyzed RNA-Seq data, H3K27ac and H3K4me1 chromatin immunoprecipitation-sequencing (ChIP-Seq) data, and previously published chromatin interaction analysis by paired-end tag sequencing (ChIA-PET) data to characterize the chromatin landscapes within the know JIA-associated risk loci. In both neutrophils and primary CD4+ T cells, the majority of the JIA-associated LD blocks contained H3K27ac and/or H3K4me1 marks. These LD blocks were also binding sites for a small group of transcription factors, particularly in neutrophils. Furthermore, these regions showed abundant intronic and intergenic transcription in neutrophils. In neutrophils, none of the genes that were differentially expressed between untreated JIA patients and healthy children was located within the JIA risk LD blocks. In CD4+ T cells, multiple genes, including HLA-DQA1, HLA-DQB2, TRAF1, and IRF1 were associated with the long-distance interacting regions within the LD regions as determined from ChIA-PET data. These findings suggest that aberrant transcriptional control is the underlying pathogenic mechanism in JIA. Furthermore, these findings demonstrate the challenges of identifying the actual causal variants within complex genomic/chromatin landscapes.

## Introduction

In recent years, there has been considerable progress in our ability to identify genetic risk in juvenile idiopathic arthritis (JIA) [1, 2]. Furthermore, we are beginning to understand the contribution that genetic variance makes to the heterogeneity of clinical response to first-line agents such as methotrextate [3]. However, taking genetic risk information and integrating it into a coherent, testable hypothesis of disease pathogenesis has remained challenging. A recent genetic fine mapping study by Hinks et al [4] illustrates that challenge. While these authors were able to identify multiple previously unknown regions of genetic risk for JIA using the Illumina Immunochip, the majority of the risk-associated variants were located within non-coding regions of the genome. In this regard, JIA resembles almost every other complex trait that has been studied using genome-wide association studies (GWAS) [5, 6], including immunologic/autoimmune diseases [7]. Thus, although it’s common to identify genetic risk loci in terms of the closest protein-coding gene, we are learning that the genome is far more complex than previously imagined, and that the presence of a risk-conferring single nucleotide polymorphism (SNP) near a specific gene is not *prima facie* evidence that the causal variants have anything to do with that particular gene [8].

We have previously demonstrated how the field might begin to make sense of the wealth of genetic data that is being generated via GWAS and fine mapping studies [9]. We recently demonstrated that the majority of the disease-associated SNPs identified on a genetic fine mapping study by Hinks et al that are situated within the non-coding genome are situated within linkage disequilibrium (LD) blocks that are enriched for H3K4me1/H3K27ac histone marks, epigenetic signatures associated with enhancer function, in both neutrophils and CD4+ T cells. Several of these same LD blocks contain non-coding RNAs that were identified on RNA sequencing (RNA-Seq) and verified by reverse transcriptase polymerase chain reaction (rtPCR) [9].

Our earlier paper focused entirely on the novel risk regions identified in the Hinks study [4]. In the current study, we examined additional regions of genetic risk as recently reviewed by Hersh et al [1] and Herlin et al [2]. We demonstrate how understanding the transcriptome and functional, non-coding genome of allows us to better understand the nature of genetic risk in JIA. In the current paper, we examine the chromatin landscapes in both CD4+ T cells and neutrophils. The former are widely accepted as important mediators of the pathobiology of JIA [10], and the latter have become the subject of increasing interest from the standpoint of the role(s) in childhood onset rheumatic diseases [11]. We have previously shown, for example that neutrophils from children with JIA display specific aberrations in gene expression that are linked to perturbations in fundamental metabolic processes [12]. We used publically available genomic data as well as our own RNA sequencing data to gain mechanistic insights into JIA disease processes from genetic risk data.

## Materials and Methods

*Patients and Patient Samples*–Neutrophils for RNA sequencing (RNA-Seq) were obtained from 3 children (2 girls and 1 boys) with newly diagnosed, active, untreated JIA (ADU group) and 2 healthy control children (HC group) recruited from the University of Oklahoma General Pediatrics clinic. Patients with JIA all had the polyarticular, rheumatoid factor negative phenotype as defined by the International League Against Rheumatism [13]. In addition, we studied pediatric-specific transcriptomes in CD4+ T cells from 5 healthy children. For the CD4+ T cell sample collection, children were excluded if they were taking antibiotics or steroids, had fever within the previous 36 hr, had an underlying autoimmune (e.g., type 1 diabetes) or inflammatory disease (e.g., asthma), or had a body mass index of > 30.

Whole blood was drawn into 10 mL CPT tubes (Becton Dickinson, Franklin Lakes, NJ), which is an evacuated blood collection tube system containing sodium citrate anticoagulant and blood separation media composed of a thixotropic polyester gel and a FICOLL^TM^ Hypaque^TM^ solution. Cell separation procedures were started within one hour from the time the specimens were drawn. Neutrophils were separated by density-gradient centrifugation at 1,700x g for 20 minutes. After removing red cells from neutrophils by hypotonic lysis, neutrophils were then immediately placed in TRIzol^®^ reagent (Invitrogen, Carlsbad, CA) and stored at −80°C until used for RNA isolation. Cells prepared in this fashion are more than 98% CD66b+ by flow cytometry and contain no contaminating CD14+ cells, as previously reported [14]. Thus, although these cell preparations contained small numbers of other granulocytes, they will be referred to here as “neutrophils” for brevity and convenience.

CD4+ T cells were obtained using the same collection techniques. CD4+ T cells were isolated from PBMC using a negative selection strategy.

All human subjects research was reviewed and approved by the University of Oklahoma and the University at Buffalo Woman & Children’s Institutional Review Boards (IRB). All research was conducted in accordance with the IRB-approved protocols. Written informed consent was obtained from the parents of children (patients and controls) participating in this study.

### RNA isolation and sequencing

Total RNA was extracted using Trizol^®^ reagent according to manufacturer's directions. RNA was further purified using RNeasy MiniElute Cleanup kit including a DNase digest according to the manufacturer's instructions (QIAGEN, Valencia, CA). RNA was quantified spectrophotometrically (Nanodrop, Thermo Scientific, Wilmington, DE) and assessed for quality by capillary gel electrophoresis (Agilent 2100 Bioanalyzer; Agilent Technologies, Inc., Palo Alto, CA). Paired-end cDNA libraries were prepared for each sample and sequenced using the Illumina TruSeq RNA Sample Preparation Kit by following the manufacture's recommended procedures and sequenced using the Illumina HiSeq 2500. Library construction and RNA sequencing were performed in the University at Buffalo Genomics and Sequencing Core Facility.

### Analysis of RNA-Seq Data

The raw reads obtained from paired-end RNA-Seq were mapped to human genome hg19, downloaded from the University of California Santa Cruz Genome Bioinformatics Site (http://genome.ucsc.edu), with no more than 2 read mismatches using tophat v2.0.10 [15]. Gene expression level was calculated as FPKM (Fragments Per Kb per Million reads) with Cufflinks v2.2.1 [16], the annotation gtf file provided for genes is gencode v19 annotation gtf file from GENCODE (http://www.gencodegenes.org) [17]. We used Cuffdiff v2.2.1 [18] for pairwise comparisons of ADU and HC to identify differentially expressed genes in human neutrophils, with a false discovery rate of 5%.

### Verification of non-coding RNA transcripts identified in neutrophils within the JIA-Associated LD blocks

Complementary DNA was synthesized from total RNA using an iScript cDNA Synthesis kit (Bio-Rad). RT-PCR was performed using a HotStar Master PCR kit (Qiagen) with a Veriti thermocycler (Life Technologies) and included a control with no reverse transcription to exclude the possibility of an artifactual result due to contaminating genomic DNA. The temperature profile consisted of an initial step of 95°C for 10 minutes, followed by 35 cycles of 95°C for 30 seconds, 60°C for 30 seconds and 72°C for 30 seconds, and then a final step 72°C for 10 minutes. PCR products were resolved in 1.5% agarose gel. The nucleotide sequences of the primers were as follows: for ncTNFa (chr6: 31,544,379-31,544,485), 5′-CCTAATTCTGGGTTTGGGTTTGGG-3′ (forward) and 5′-CCTACTTTCACCTCCATCCATCCT-3′ (reverse); for ncSTAT4 (chr2:191,915,516-191,915,617), 5′-GTCTTGTGCAACTTCTTCCTTTC-3′ (forward) and 5′-ACCCTGTGACTGTTTGAGATTAC-3′ (reverse).

### Defining LD regions

SNPs used in this query are listed in Table 1 and were previously reviewed by Hersh et al [1], Herlin et al [2] and Hinks et al [19]. We used a SNP Annotation And Proxy search (SNAP) database (http://www.broadinstitute.org/mpg/snap) [20] to define LD blocks based on the location of each SNP. In brief, we used the settings as follows: SNP dataset: 1000 Genome pilot 1 and HapMap 3 (release 2), r^2^ threshhold: 0.9, Population Panel: CEU, Distance limit: 500. We selected the smallest number as our start location and the largest number as our stop location for each defined LD block.

**Table 1:** Histone marks in the SNP LD blocks in neutrophils and CD4+T cells

| LD region | SNP | Neutrophils |  | CD4+T cells |  |
| --- | --- | --- | --- | --- | --- |
|  |  | H3K4me1<br>mark | H3K27ac<br>mark | H3K4me1<br>mark | H3K27ac<br>mark |
| chr1: 206940310 - 206947167 | rs1800896 | Y | N | Y | Y |
| chr1: 25197155 - 25203390 | rs4648881 | Y | N | Y | N |
| chr11: 36336263 - 36371757 | rs4755450, rs7127214 | N | N | N | N |
| chr12: 111884608 - 111932800 | rs3184504 | Y | Y | Y | Y |
| chr12: 112486818 - 112906415 | rs17696736 | Y | Y | Y | Y |
| chr13: 40299842 - 40368601 | rs7993214, rs9532434 | N | N | Y | N |
| chr13: 43056036 - 43066523 | rs34132030 | N | N | N | N |
| chr16: 11400900 - 11435990 | rs66718203 | Y | Y | Y | N |
| chr18: 12821903 - 12880206 | rs7234029 | Y | Y | Y | Y |
| chr2: 100806514 - 100837567 | rs10194635, rs6740838,<br>rs1160542 | Y | N | Y | Y |
| chr2: 191900449 - 191935804 | rs3821236 | Y | Y | Y | Y |
| chr2: 191943742 - 191970120 | rs7574865 | N | N | Y | N |
| chr22: 24234493 - 24237862 | rs755622 | Y | Y | Y | Y |
| chr3: 119125202 - 119243934 | rs4688013, rs4688011 | Y | Y | Y | Y |
| chr3: 46253650 - 46350716 | rs79893749 | Y | Y | Y | Y |
| chr4: 123141054 - 123548068 | rs17388568 | Y | Y | Y | Y |
| chr6: 112359543 - 112448654 | rs2280153 | Y | Y | Y | Y |
| chr6: 137959235 - 138006504 | rs6920220 | N | N | Y | N |
| chr6: 31492453 - 31543031 | rs1800629 | N | Y | Y | Y |
| chr7: 22774437 - 22811384 | rs6946509, rs7808122 | N | N | N | N |
| chr7: 28152193 - 28243473 | rs10280937,<br>rs73300638 | Y | Y | Y | Y |
| chr9: 123636121 - 123723351 | rs10818488 | Y | Y | Y | Y |
| chr9: 123640500 - 123706382 | rs2900180 | Y | Y | Y | Y |

Table 1: Histone marks in the SNP LD blocks in neutrophil and CD4+T cells
| LD region | SNP | Neutrophil |  | CD4+T |  |
| --- | --- | --- | --- | --- | --- |
|  |  | H3K4me1 mark | H3K27ac mark | H3K4me1 mark | H3K27ac mark |
| chr1: 206940310 - 206947167 | rs1800896 | Y | N | Y | Y |
| chr1: 25197155 - 25203390 | rs4648881 | Y | N | Y | N |
| chr11: 36336263 - 36371757 | rs4755450, rs7127214 | N | N | N | N |
| chr12: 111884608 - 111932800 | rs3184504 | Y | Y | Y | Y |
| chr12: 112486818 - 112906415 | rs17696736 | Y | Y | Y | Y |
| chr13: 40299842 - 40368601 | rs7993214, rs9532434 | N | N | Y | N |
| chr13: 43056036 - 43066523 | rs34132030 | N | N | N | N |
| chr16: 11400900 - 11435990 | rs66718203 | Y | Y | Y | N |
| chr18: 12821903 - 12880206 | rs7234029 | Y | Y | Y | Y |
| chr2: 100806514 - 100837567 | rs10194635, rs6740838, rs1160542 | Y | N | Y | Y |
| chr2: 191900449 - 191935804 | rs3821236 | Y | Y | Y | Y |
| chr2: 191943742 - 191970120 | rs7574865 | N | N | Y | N |
| chr22: 24234493 - 24237862 | rs755622 | Y | Y | Y | Y |
| chr3: 119125202 - 119243934 | rs4688013, rs4688011 | Y | Y | Y | Y |
| chr3: 46253650 - 46350716 | rs79893749 | Y | Y | Y | Y |
| chr4: 123141054 - 123548068 | rs17388568 | Y | Y | Y | Y |
| chr6: 112359543 - 112448654 | rs2280153 | Y | Y | Y | Y |
| chr6: 137959235 - 138006504 | rs6920220 | N | N | Y | N |
| chr6: 31492453 - 31543031 | rs1800629 | N | Y | Y | Y |
| chr7: 22774437 - 22811384 | rs6946509, rs7808122 | N | N | N | N |
| chr7: 28152193 - 28243473 | rs10280937, rs73300638 | Y | Y | Y | Y |
| chr9: 123636121 - 123723351 | rs10818488 | Y | Y | Y | Y |
| chr9: 123640500 - 123706382 | rs2900180 | Y | Y | Y | Y |

### ENCODE Transcription Factor Binding Site (TFBS) enrichment

ENCODE TFBS data were downloaded from UCSC Genome Browser ENCODE data (http://hgdownload.cse.ucsc.edu/goldenPath/hg19/encodeDCC/wgEncodeRegTfbsClustered/). Only the TFBS information annotated for blood cells was used. The whole genome was binned to 100bp bins and intersected with H3K4me1 or H3K27ac peak regions, which were used as background. Fisher’s exact test was applied to test the significant of the enrichment of TFBS of each transcription factor within H3K4me1 or H3K27ac peaks in LD blocks compared to peak regions in non-LD blocks across the whole genome. We set the cutoff of false discovery rate (FDR) to 0.05 for all the analyses.

### Identification of differentially expressed neutrophil genes within LD regions

To determine whether any of the differentially expressed genes (DEGs) identified in the comparison of neutrophil transcriptomes of children with untreated JIA (ADU) and HC were located within the JIA-associated LD blocks, we used gencode v19 annotation and intersectBed to refine the gencode data to focus specifically on those LD blocks.

### Identification of H3K4me1/H327ac histone marks within LD regions

We used H3K4me1/H3K27ac chromatin immunoprecipitation-sequencing (ChIP-Seq) data previously generated by our group from neutrophils of healthy adults [9] as our reference source for H3K4me1/H3K27ac genomic locations in neutrophils (GEO accession # GSE66896). The H3K4me1/H3K27ac ChIP-Seq data reported on the Roadmap Epigenomics collection were used as our reference source for H3K4me1/H3K27ac genomic locations in CD4+ T cells (GEO accession # 198927). The intersection of H3K4me1/H3K27ac peaks within LD regions was obtained using the bedtools intersect procedure [21] as described in our previous paper [9].

### Identifying chromatin interactions in CD4+ T cells within JIA risk loci

Functional DNA elements such as enhancers do not always regulate the nearest gene. Indeed, in a recent study of chromatin conformation using HiC in T and B cell lines, Martin et al [8] showed that > 80% of long distance chromatin interactions occurred at distances of > 500 kb. We therefore queried an existing chromatin interaction analysis by paired-end tag sequencing (CHIA-PET) dataset for CD4+ T cells (GSE32677) [22] to determine chromatin interactions within the JIA-associated risk loci. This data set used H3K4me2 as a marker for active enhancers, and thus provides a means of estimating those genes/gene networks that are regulated by specific enhancers or groups of enhancers. The chromosome coordinates for the observed 6520 long-distance chromatin interactions were converted from hg18 to hg19 using UCSC Genome browser LiftOver tool (http://www.genome.ucsc.edu/cgi-bin/hgLiftOver). For these analyses, we queried all known JIA-associated risk regions.

### Identifying molecular pathways of disease-associated SNPs

One means of using genetic risk data to provide mechanistic insights is to identify specific biological pathways encompassed by SNP-associated genes [23, 24]. However, there is a considerable breadth of opinion about what constitutes a “SNP-associated gene,” as the actual causal variants may be a considerable distance (in genomic terms) from the risk-associated SNP. We therefore used the method of Brodie et al [25] to identify molecular pathways encompassed by JIA-associated SNPs across broad genomic distances. We queried regions 200 kb upstream and downstream of the JIA-associated SNPs, as outlined in [25] to define genes and pathways associated with JIA risk SNPs.

### Data availability

The authors state that all data necessary for confirming the conclusions presented in the article are represented fully within the article.

## RESULTS

### Results from RNA-Seq and location of differentially expressed genes within LD blocks

Our previous work had queried new SNPs recently identified on a genetic fine mapping study published by Hinks et al [4]. In the current study, we queried SNPs that were identified in reviews by Hersh et al [1] and Herlin et al [2] and those SNPs in Hinks et al [19] that have not been analyzed in previous work and have established associations with JIA (Table 1). We used these SNPs to generate linkage disequilibrium blocks (LD blocks) using SNAP [20] and used these defined regions to determine DEGs located within these regions. We included SNPs located within non-coding regions of the genome (introns or intergenic areas), i.e., the majority of the JIA-associated SNPs.

We identified 99 genes that showed differential expression between children with newly diagnosed, untreated JIA (ADU) and HC (FDR < 0.05). Predictably, GO analysis of these genes (compared to the background of neutrophil-expressed genes) showed enrichment for responses to viruses and as well as other immune responses. None of the differentially expressed genes was located within the LD blocks defined by the JIA-associated SNPs. This finding is consistent with what we have seen in whole blood microarray expression profiles [26].

### Verification of non-coding transcripts within neutrophils in JIA-associated LD blocks

Review of RNA-Seq data using the University of California Santa Cruz (UCSC) Genome Browser suggested the presence of multiple non-coding RNA transcripts in neutrophils within the JIA-associated LD blocks. We used an rtPCR approach to confirm the presence of 2 of these transcripts, i.e., those within the LD blocks containing the rs3821236 and rs1800629 SNPs. Figure 1 shows detectable expression of these transcripts in neutrophils from a healthy child and a JIA patient. This finding corroborates the idea that the JIA-associated LD blocks contain functional, non-coding RNA transcripts.

**Figure 1.**
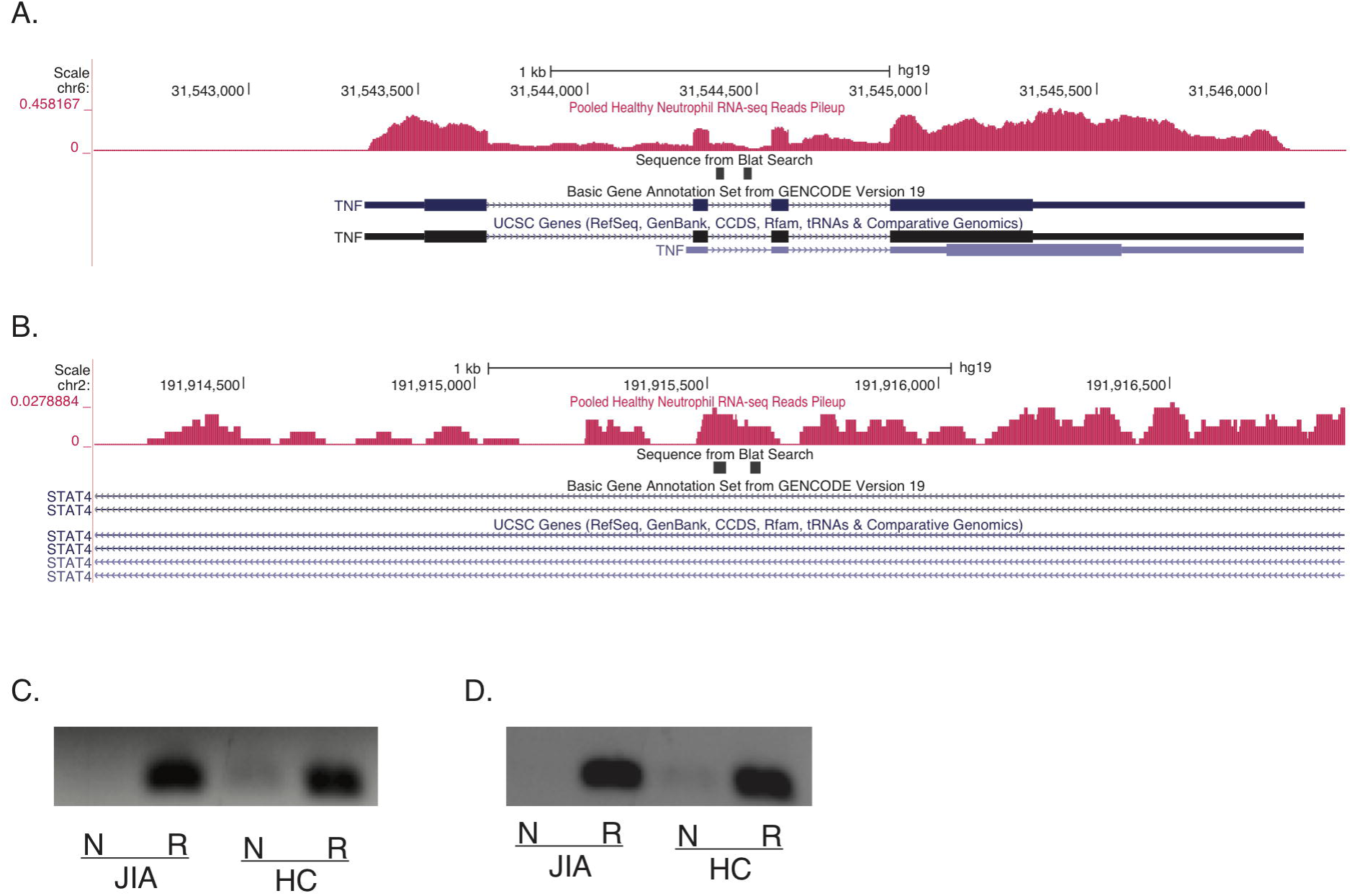
UCSC Genome Browser screen shot showing abundant intronic transcription in the *TNF* gene (**A**) and in the STAT4 gene (**B**) within introns. The black bars indicate the region amplified by PCR experiments, as described in the *Methods* section. **c** and **d**, Agarose gel images showing qualitative PCR amplification of intronic transcription (ncRNA) in the *TNF* gene (**C**) and in the STAT4 gene (**D**) in neutrophils. N = neutrophil RNA not subjected to reverse transcription; R = neutrophil RNA subjected to reverse transcription prior to amplification. Neutrophils were isolated from patients with juvenile idiopathic arthritis (JIA) and healthy children (HC). Absence of signal in PCR amplifications that were performed without prior reverse transcription indicates that the signal is not being produced by contaminating DNA.

### Location of H3K4me1/H3K27ac marks within LD blocks

For these analyses, we examined only those LD blocks not previously examined in our earlier paper [9] as noted in the Methods section. We identified LD blocks for all of the 30 risk SNPs, which were located in a total of 23 LD blocks. For those 23 LD blocks, we examined whether there are functional elements located in neutrophils or CD4+ T cells within these risk-conferring regions.

Our first analyses were in neutrophils. Of the 23 risk loci, 16 LD blocks contained H3K4me1 marks, while 14 LD blocks contained H3K27ac marks; 13 LD blocks contained both H3K4me1 and H3K27ac marks (Table 1). In all, 17 out of the 23 LD blocks contained either H3K4me1 or H3K27ac marks. The H3K4me1 and H3K27ac marks are significantly enriched in these LD block regions compared with non-LD block regions as determined by Fisher’s exact test (p-value < 2.2e-16 for both H3K4me1 and H3K27ac marks). Representative screen shots from the UCSC genome browser within 2 of the LD regions of interest are shown in Figure 2. The LD block containing rs755622, for example (Figure 2A), includes the promoter for the macrophage inhibitory factor (MIF) gene, but this region is also rich in TF binding regions and contains H3K4me1/H3K27ac histone marks. Elevated MIF levels have been found in patients with all 3 major JIA subtypes [27], although these levels are most elevated in children with the oligoarticular and systemic forms of the disease. Similarly, the neutrophil chromatin landscape around rs2900180 (Figure 2B) shows considerable genomic complexity. There is an expressed gene within this region, i.e., the tumor necrosis factor receptor-associated factor 1 [TRFA1]); there has been strong interest in the tumor necrosis factor (TNF) signaling pathways since the efficacy of anti-TNF therapies was demonstrated in children with JIA [28]. At the same time, this region contains both H3K4me1/H3K27 marked enhancers and abundant TF binding not isolated to the TRAF promoter. Similarly, **Figure S1** shows multiple functional elements adjacent to potentially disease-relevant genes such as PTPN2 (**S1A**) and NFkB inhibitor-like protein 1 (NFkBIL1, **S1B**). Thus, these findings corroborate what we previously reported [9] regarding the genetic risk regions identified in the Hinks paper [4], i.e., the genetic risk largely encompasses regions of the genome rich in functional elements that serve to regulate and coordinate transcription on a genome-wide basis.

**Figure 2.**
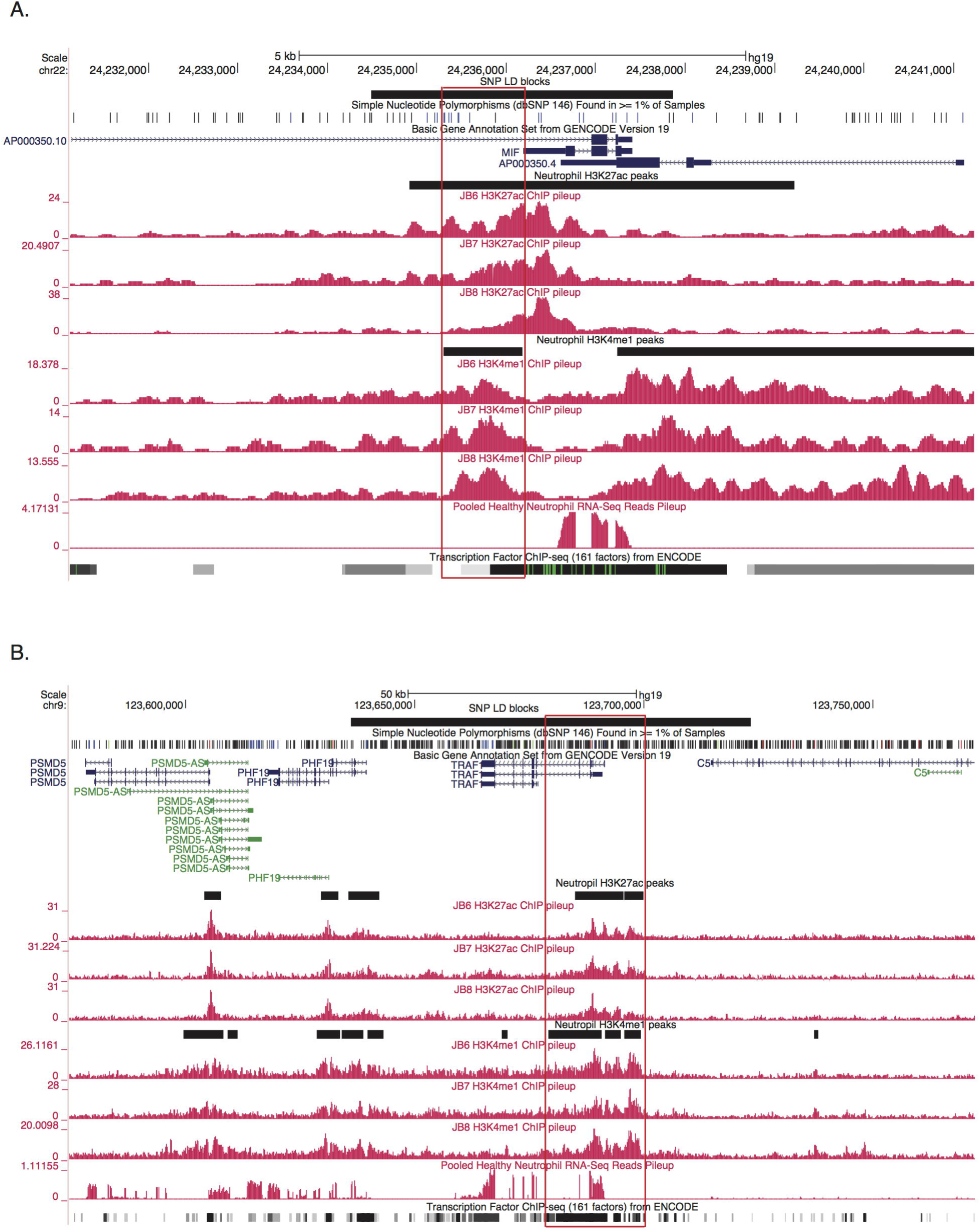
Representative screen shots from the UCSC Genome Browser showing functional elements within neutrophil genomes. The black horizontal bar at the top represents the LD blocks of the corresponding GWAS SNP. The black horizontal bars between individual tracks represent H3K27ac and H3K4me1 peak regions generated from ChIP-Seq data. The grey bars at the bottom of each image represent the transcription factor ChIP-seq of 161 factors from ENCODE with factorbook motifs. Panel **(A)** shows the LD block of rs755622 in neutrophils. Note there are expressed genes within this LD block, and the LD block contains H3K27ac and H3K4me1 regions with multiple potential TFBS within those regions. Panel **(B)** shows the LD block of rs2900180 in neutrophils. Note that there is one gene, the tumor necrosis factor receptor associated factor (TRAF1), which is expressed within this LD block. This LD block also contains both H3K27ac and H3K4me1 peak regions with multiple TFBS within those regions. Potential active enhancers (region containing both H3K27ac and H3K4me1 peak regions) are highlighted in red.

We next examined H3K4me1/H3K27ac marked regions in CD4+ T cells in those same 23 LD blocks. We queried whether there are functional elements located in CD4+ T cells within these risk-conferring regions. Of the 23 LD blocks, 20 contained H3K4me1 marks and 15 contained H3K27ac marks. All of the 15 LD blocks that contained H3K27ac marks also contained H3K4me1 marks (Table 1). Thus, 20 out of 23 of the LD blocks contained either H3K27ac or H3K4me1 marks in CD4+ T cells. The H3K27ac and H3K4me1 marks are significantly enriched in these LD block regions compared with non-LD block regions as determined by Fisher’s exact test (p-value < 2.2e-16 for both H3K27ac and H3K4me1 marks).

### Transcription factor enrichment within the LD blocks

While H3K4me1/H3K27ac marks in and of themselves do not unambiguously identify functional enhancers[29], the appearance of transcription factor binding sites within H3K4me1/H3K27ac-marked regions is strong supportive evidence that these regions are functional [30]. Therefore, for the each of the 38 LD blocks containing the 51 JIA-associated risk SNPs, we queried whether there was significant enrichment for TF binding in the H3K27ac/H3K4me1 marked regions compared to non-LD block regions.

In these analyses, we included genetic risk information from the Hinks, Hersh, and Herlin papers. Thus, to determine whether there was additional evidence that the H3K4me1/H3K27ac marked regions within the JIA-associated LD blocks had functional significance, we queried whether there were significantly enriched TFBS within those loci compared to non-LD regions that have H3K4me1/H3K27ac marks.

For neutrophils, we observed highly enriched TF binding in promoter regions (defined as (-5K, 1K) of TSS) within these LD blocks. TFs that were particularly enriched included POLR2A, CHD1 and TAF1 (Figure 3A). We note that RUNX3 binding was also enriched within the H3K27ac-marked regions (Figure 3A), an interesting finding in that the RUNX3 locus itself is associated with JIA disease risk. This locus is characterized, in neutrophils, by dense intergenic TF binding, the presence of both H3K4me1 and H3K27ac peaks overlapping with the TF binding regions, and both intronic and intergenic RNA transcripts (not yet verified by rtPCR experiments; see **Figure S3**). These findings suggest complex relationships between enhancers and promoters, and perhaps non-coding RNA transcripts, leading to complex levels of gene regulation in neutrophils in health and as well as in disease states such as JIA.

**Figure 3.**
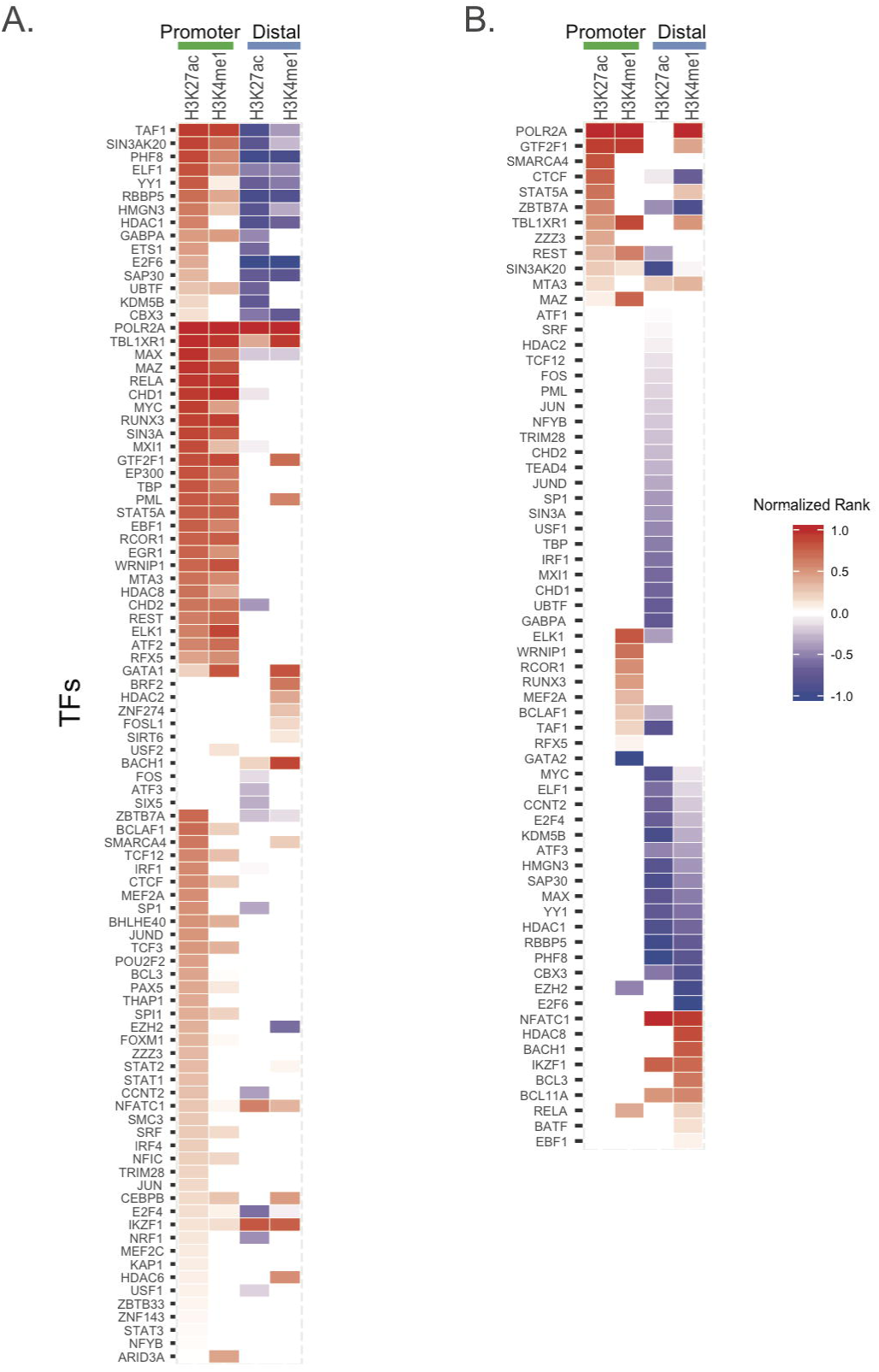
Enrichment of TFs in H3K4me1/H3K27ac mark regions. Heatmap of significantly enriched TFs with their TFBS in H3K27ac or H3K4me1 peak regions in **(A)** neutrophil cells and **(B)** CD4+T cells, considering promoter and distal regions separately. TFs are ranked according to FDR (the one lowest FDR with the highest rank) and normalized to (0,1) by dividing the absolute highest rank value for enrichment (red) and depletion (blue) separately. (normalized rank: > 0 enrichment; <0 depletion).

For CD4+ T cells, we observed a limited number of TFBS that were significantly enriched in the LD blocks conferring risk for JIA that co-localized with H3K27ac/H3K4me1 histone marks (Figure 3B). These regions were seen almost exclusively in promoters, defined as −5K − 1K of the transcription start site (TSS). These transcription factors included POLR2A and GTF2F1. The majority of the distal (non-promoter) regions within these LD blocks were relatively depleted of TFBS. Thus, the LD blocks with H3K27ac/H3K4me1 marks are not highly enriched with TFs. This finding contrasts with what we observed in neutrophils, as noted above, where there is significant enrichment of TF binding within H3K4me1/H3K27ac-marked regions.

### Chromatin interactions in JIA risk loci

We used the 6520 long-distance chromatin interactions from a previously-reported ChIA-PET data analysis of CD4+ T cells [22] to define chromatin interactions in all the JIA-associated LD blocks. We identified 11 LD blocks that contained 48 long-range interactions. We note that all 6 of the interaction regions within the LD block containing the SNP, rs10818488, are identical with the regions within the LD block of rs2900180. This left 42 unique long-range interactions, 11 of which displayed more than one interaction within or intersected with the same LD blocks (**S. Table 1**). For these 42 long-range interactions, 20 were promoter-promoter (defined as (-5K, 1K) of TSS) interactions and 21 were promoter-distal (non-promoter) interactions, and one was a distal-distal interaction.

We next analyzed the CHIA-PET data in light of transcriptomes from CD4+ T cells derived from healthy children. We identified several genes within the CHIA-PET interacting regions, including ILB, DDX39B, RASSF5 and JAK1, that demonstrated high levels of expression (average FPKM > 100). We note that there are multiple small RNA molecules (snoRNA, miRNA, misc_RNA, snRNA) that are encoded within these regions, but the sequencing libraries were not prepared to detect their presence since the FPKM of those molecules is 0.

Among the 42 long-range interactions, 66 regions contained both H3K4me1 and H3K27ac marks, while 2 regions contained only H3K27ac marks and 11 contained only H3K4me1 marks. Only 5 of the interacting regions contained neither H3K4me1 nor H3K27ac marks. There were in all 61 genes associated with these long-range interacting regions. The genes associated with regions containing both H3K4me1 and H3K27ac marks have average higher FPKM than those genes that contained only H3K4me1 marks or those with no histone marks (**Figure S2**).

The CHIA-PET data point to complex layers of interaction and gene regulation within the JIA-associated risk loci. Note, for example, multiple points of interaction between the IRF1 and Corf56 genes, a locus that also encodes a Y-RNA species. These genes are physically adjacent to one-another on chromosome 5, as shown in Figure 4A. This region is dense with functional elements, including an H3K4me1/H3K27ac-marked enhancer, dense transcription factor binding, and multiple DNase1 hypersensitivity sites. It is interesting to note that our previous work from whole blood expression profiles of untreated JIA patients has suggested an important role for IRF1 networks in JIA pathogenesis [26]. Equally interesting is the detected physical interaction between the HLADQB1 and HLADQB2 loci. Association between JIA risk and class II HLA alleles has been long recognized, although the alleles at the DRB1 locus are thought to have stronger effects than those at the DQB1 or DQB2 loci [31]. The genome browser shot in Figure 4B demonstrates the complexity of this locus. Complex interactions between TRAF1 and multiple adjacent genes (C5, PHF19, and CNTRL) as well as TRAF1 intragenic interactions were also observed on the CHIA-PET data by Chepelev et al [22].

**Figure 4.**
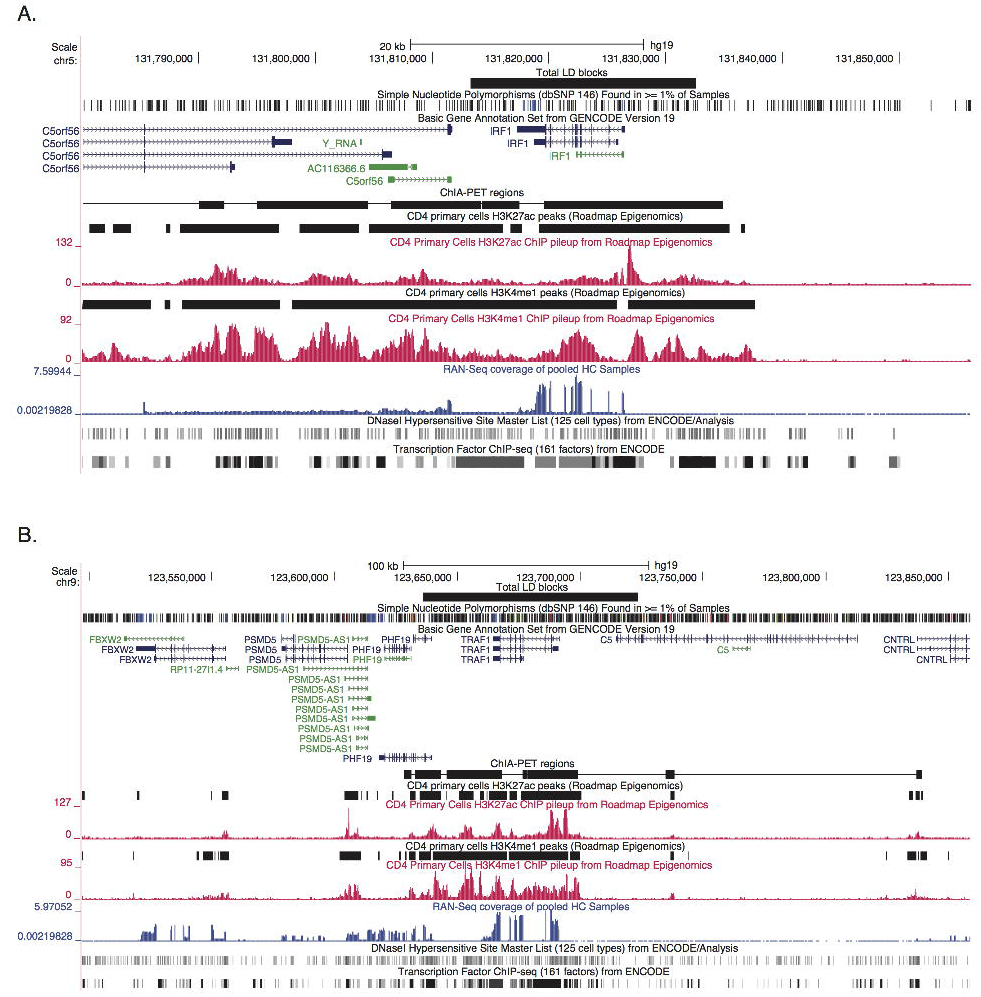
Genome browser shot of the IRF1/C5orf56 locus in CD4 primary T cells **(A)** which is an epigenetically rich region showing dense transcription factor binding, multiple DNase1 hypersensity sites, and active enhancers with both H3K27ac and H3K4me1 marks. **(B)** TRAF1 locus with rich epigenetic architecture in this locus, including multiple TF binding sites, DNase1 hypersenitive sites, and active enhancers (H3K27ac and H3K4me1 marked regions).

### Enriched molecular pathways of disease-associated SNPs

We used the method of Brodie et al [25] and the PANTHER database [32] to identify significantly enriched biological process and pathways for the 246 protein-coding genes and processed transcripts associated with the 51 JIA risk SNPs. Using a cutoff of Bonferroni <=0.05, the enriched PANTHER pathways are related to interleukin signaling and inflammation mediated by chemokine and cytokine signaling. The enriched biological processes involve antigen processing and presentation, cellular defense response as well as other immunologically relevant processes (**S. Table 2**).

## Discussion

In this paper, we mined existing data sets in order to gain a better understanding of the nature of genetic risk in JIA. We find that most of the JIA-associated SNPs lie within the non-coding genome, in regions that are dense with TF binding sites, DNaseI hypersensitive sites, epigenetic signatures of enhancer function, and, in the case of neutrophils, non-coding RNA transcripts. These findings demonstrate the importance of considering the chromatin context around disease-associated SNPs, which may have no functional association at all with the “nearest gene.”

At the completion of the Human Genome Project, there was some surprise and even a little dismay at how little of the genome contains protein-encoding genes. When the entire project was done, only ~2% of the DNA was found to ENCODE proteins [33]. One of the transformative findings from the ENCODE and Roadmap Epigenomics projects has been the discovery of the richness of the non-coding genome. Enhancers, insulators, CTCF binding sites (which regulate chromatin conformation) etc are broadly distributed throughout the genome and demonstrate that the processes of regulating gene expression and coordinating expression on a genome-wide basis is extraordinarily complex [34]. The field of immunology is now making considerable progress in identifying these functional elements and how they regulate transcription and therefore immune responses [35].

The findings from ENCODE and Roadmap Epigenomics, and the work we describe here, demonstrate that identifying specific causal variants in JIA is going to be an ongoing challenge. A primary reason for this is the complexity of the risk loci. While many of these loci encompass genes that are of strong immunologic interest (e.g., MIF, TRAF1, PTNP2, NFkBIL1), these same loci are also rich in other functional elements that regulate transcription on a global basis. For example, Figure 2A is a simplified genome browser screen shot that shows the neutrophil chromatin landscape around rs755622.

While the actual SNP is within the promoter of the macrophage inhibitory factor gene, the entire LD block contains both H3K4me1 and H3K27ac histone marks (strong evidence for an active enhancer in this region) as well as rich TF binding that is not localized exclusively to the promoter. Similar genomic features can also be seen in this region in CD4+ T cells (data not shown). We also find find a large degree of complexity around rs2900180 (Figure 2B), and rs2847293/rs7234029, which are in the same LD block. The region containing rs2847293/rs7234029 is of broad general interest to the field of immunology, as this LD block includes the *PTPN2* gene. While PTPN2 is generally considered a regulator of T cell responses, this protein tyrosine phosphatase is broadly expressed in innate immune cells, including in neutrophils, where it serves as a negative regulator of cytokine signaling [36]. However, the PTPN2 locus is also characterized by H3K4me1/H3K27ac-marked enhancers in neutrophils and CD4+ T cells, as well as an apparent non-coding RNA molecule in the intron between the first and second exons in neutrophils (**Figure S1A**). Thus, while it is tempting to focus on the protein-encoding genes within these regions as the location of the causal variants that lead to perturbed immune function, our data invite other interpretations. Enhancers, for example, may function at considerable distances (in genomic terms) from the genes they regulate, and often do not regulate the closest gene [37, 38]. This idea is shown in the recent work of Martin et al [8]. Using Hi-C chromatin conformation capture approaches, these authors demonstrated that > 80% of long-distance chromatin interactions within T and B cell lines occurred at distances > 500 MB. Thus, until there is more solid experimental evidence to indicate otherwise, we must be open to the possibility that it is the enhancer function of these regions and/or the function of the ncRNA that are perturbed by genetic variance rather than specific functions or regulation of the nearby proteins.

This idea is corroborated by our analysis of the published CD4+ T cells CHIA-PET data set. CHIA-PET, uses cross-linked DNA to identify physical interactions between regions of DNA, which may either be in close proximity or many megabases apart on a specific chromosome (or even regions that are on different chromosomes) [22, 39]. The dataset we queried used H3K4me2 as a general mark for active enhancers. We found that the JIA-associated genetic risk loci display abundant and complex physical interactions with (usually) nearby genomic elements. The region encoding the TRAF1 gene is emblematic of this complexity. We detected interactions between an H3K4me2-marked region adjacent to the TRAF1 gene and at least 3 other genes. These genes included C5, the 5^th^ component of the complement cascade, an important inflammatory mediator. The region also interacts with PHF19, which is itself a regulator of gene expression. PHF19 is a member of the polycomb repressive complex-2 (PRC-2) that is involved in gene silencing [40]. The significance of the interaction between the TRAF1 region and CNTRL (formerly for as CEP110), a centrosomic protein [41] is not immediately clear.

ENCODE also revealed that much of the genome is transcribed, and that many (if not most) of these non-coding RNA species are functional. In rheumatology, miRNA are the best-known of these non-coding RNA families, as numerous papers have shown links between specific miRNA molecules and specific immunologic features of the rheumatic diseases (e.g., [42]). Our own work has recently shown extensive re-wiring of mRNA-miRNA networks within JIA neutrophils [14]. However, the non-coding RNA world is broad, and includes RNA species that regulate chromatin accessibility [43], serve enhancer functions [44-46] and encode for small peptides [47]. Our current work also demonstrates the presence of at least two intronic, non-coding RNA molecules in the JIA-associated genetic risk loci. Such transcripts are abundant in neutrophils [48] and may regulate the decay of otherwise unstable mRNA transcripts [45]. The specific roles of these transcripts in neutrophil biology and/or the pathogenesis of JIA remain to be clarified. Their presence, however, serves to demonstrate that the localization of a disease-associated polymorphism within a specific gene does not necessarily mean that genetic risk is conferred by alterations in the protein-coding function of that specific gene.

This paper corroborates and expands upon a recent report from our laboratory demonstrating that regions of genetic risk in JIA generally do not encode for genes that show differential expression when children with the disease are compared with healthy children [26]. Using whole blood gene expression data from subjects of a National Institutes of Health-funded clinical trial, corroborated by an independent patient cohort, we reported that none of the >150 genes that showed differential expression between patients and healthy controls were encoded within the LD blocks associated with JIA risk. Thus, the mechanisms through which genetic variance leads to perturbed gene expression are likely to be more complex than was previously considered.

This paper adds to the mounting evidence that suggests that the field of pediatric rheumatology may require a fundamental re-assessment regarding theories of JIA pathogenesis. Long accepted as an “autoimmune disease” triggered by the recognition of a “self” peptide by the adaptive immune system, new evidence from the fields of genetics and genomics ought to prompt a reconsideration of this view. We propose, instead, that JIA emerges because of genetic and epigenetically-mediated alterations in leukocyte genomes. These alterations, we propose, involve both innate and adaptive immune systems and lead to expression of inflammatory mediators in the absence of the external signals normally required to initiate or sustain an inflammatory response. Our work in both whole blood buffy coats [49] and neutrophils [12] is consistent with that interpretation of the available gene expression and genetic data.

In conclusion, we demonstrate that the known genetic risk loci for JIA are enriched for multiple functional elements within the non-coding genome of human neutrophils. These functional elements include rich TF binding sites, H3K4me1 and H3K27ac-marked enhancers, and non-coding RNAs. We believe that these data as well newer understandings of genetic risk and genome function invite new understanding of JIA pathogenesis and may provide the basis for radically new approaches to therapy.

## Acknowledgements

This work was supported by R01-AR060604 from the National institutes of Health (USA)

## Footnotes

Lisha Zhu and Kaiyu Jiang contributed equally to this work.

## Authors’ contributions

Lisha Zhu and Karstin Webber performed the primary bioinformatics analyses.

Kaiyu Jiang prepared the neutrophils and CD4+ T cells for RNA sequencing, directed and coordinated the RNA sequencing experiments, and performed rtPCR for non-coding transcripts.

Laiping Wong performed analyses to identify differentially expressed genes within the JIA-associated LD blocks.

Yanmin Chen assisted with cell isolation and RNA preparation for both neutrophils and CD4+ T cells.

Tal Liu oversaw and directed the bioinformatics analysis performed by Karstin Webber and Lisha Zhu.

James N. Jarvis designed the study, directed its implementation, as directed data analysis and interpretation.

## Contributor Information

Lisha Zhu

Kaiyu Jiang

Karstin Webber

Laiping Wong

Tao Liu

Yanmin Chen

James N. Jarvis

